# Human coronaviruses activate and hijack the proteostasis guardian HSF1 to enhance viral replication

**DOI:** 10.1101/2022.12.22.519205

**Authors:** Silvia Pauciullo, Anna Riccio, Antonio Rossi, Silvia Santopolo, Sara Piacentini, M. Gabriella Santoro

**Affiliations:** Department of Biology, University of Rome Tor Vergata, Rome, Italy; Institute of Translational Pharmacology, CNR, Rome, Italy

**Keywords:** AIRAP, antiviral, HCoV-229E, HCoV-OC43, HCoV-NL63, DTHIB, Heat shock response, Protein homeostasis

## Abstract

Organisms respond to proteotoxic stress by activating a cellular defense mechanism, known as the heat shock response (HSR), that triggers the expression of cytoprotective heat shock proteins (HSP) to counteract the damaging effects of proteostasis disruption. The HSR is regulated by a family of transcription factors (heat shock factors, HSFs); among six human HSFs, HSF1 acts as a proteostasis guardian regulating acute and severe stress-driven transcriptional responses. Seasonal coronaviruses HCoV-229E, HCoV-NL63, HCoV-OC43 and HCoV-HKU1 (sHCoV) are globally circulating in the human population. Although sHCoV generally cause only mild upper respiratory diseases in immunocompetent hosts, severe complications may occur in specific populations. There is no effective treatment for sHCoV infections, also due to the limited knowledge on sHCoV biology. We now show that both *Alpha*- and *Beta*-sHCoV are potent inducers of HSF1, selectively promoting HSF1 phosphorylation at serine-326 residue and nuclear translocation, and triggering a powerful HSF1-driven transcriptional response in infected cells at late stages of infection. Despite the coronavirus-mediated shut-down of the host cell translational machinery, high levels of selected canonical and non-canonical HSF1-target genes products, including HSP70, HSPA6 and the zinc-finger AN1-type domain-2a gene/AIRAP, were found in HCoV-infected cells. Interestingly, silencing experiments demonstrate that HSR activation does not merely reflect a cellular defense response to viral infection, but that sHCoV activate and hijack the HSF1-pathway for their own gain. Notably, nuclear HSF1 pools depletion via Direct-Targeted HSF1 inhibitor (DTHIB) treatment was highly effective in hindering sHCoV replication in lung cells. Altogether the results open new scenarios for the search of innovative antiviral strategies in the treatment of coronavirus infections.

## INTRODUCTION

Protein homeostasis is essential for life in eukaryotes. Organisms respond to proteotoxic stress by activating a cellular defense mechanism known as the heat shock response (HSR). Following its seminal discovery in *Drosophila* by Ferruccio Ritossa in the early 1960s (1), the HSR, also known as proteotoxic stress response, is recognized as a fundamental highly-conserved mechanism evolved from yeast to humans to protect cells from the damaging effects of proteostasis disruption by different types of insults including hyperthermia, exposure to toxins and alterations in the intracellular redox environment by triggering the expression of cytoprotective heat shock proteins (HSP) (2,3). HSPs, which include members of the HSP70 and HSP90 families, HSP27 and other proteins of the network, act as molecular chaperones that guide conformational states critical in the synthesis, folding, transport, assembly, ubiquitination and proteasomal degradation of proteins (2–4).

The HSR is regulated by a family of heat shock (HS) transcription factors (HSFs) that are expressed and maintained in an inactive state under non-stress conditions. The human genome encodes six heat shock factors: HSF1, HSF2, HSF4, HSF5, HSFX and HSFY that have different functions and exhibit different and tissue-specific patterns of expression (5). Among the different HSFs, HSF1 is considered to be the paralog responsible for regulating acute and severe proteotoxic stress-driven transcriptional responses, including exposures to elevated temperatures (6,7); HSF2 lacks intrinsic stress-responsiveness, but it acts as a fine tuner of the HSR contributing to inducible expression of HS genes through interplay with HSF1 (5–8).

HSF1 is a multi-domain transcription factor containing an amino-terminal DNA-binding domain, an adjacent multimerization domain, a central regulatory domain, a carboxyl-terminal coiled-coil domain and a transcriptional activation domain (4,5,9). HSF1 is generally found as an inert monomer in unstressed cells; inactive HSF1 monomers are retained in the cytoplasm in complex with regulatory proteins including HSP90, HSP70, HSP40 and the cytosolic chaperonin TRiC (TCP-1 ring complex) nanomachine (5,10). Upon stress sensing, HSF1 is derepressed in a stepwise process that involves trimerization, nuclear translocation, phosphorylation/sumoylation, and binding to DNA-sequences (heat shock elements, HSE). Functional HSE-sequences are characterized by an array of inverted repeats of the pentameric motif -nGAAn-, which are usually located in the proximal region of HSF1-responsive gene promoters, but may vary in sequence and geometry in different target genes (6,11,12). In human cells, beyond HS genes, HSF1-binding sites have been described in a broad repertoire of genes encoding proteins with non-chaperone function (13–15).

Due to the abundant amount of viral proteins rapidly synthesized in bulk, several viruses are known to depend on the host chaperone machinery for correct protein folding and assembly into viral components during different phases of the virus replication cycle (16). However, little is known on the ability of RNA viruses to activate HSF1.

Coronaviruses (CoV), members of the *Coronaviridae* family in the order *Nidovirales*, comprise a large number of enveloped, positive-sense single-stranded RNA viruses causing respiratory, enteric, renal and neurological diseases of varying severity in domestic and wild animals, as well as in humans (17). Coronaviruses have the largest identified RNA genomes (typically ranging from 27 to 32 kb); all CoV genomes are arranged similarly with a large replicase-transcriptase gene encoded within the 5’-end preceding structural proteins encoded in the 3’-end, with an invariant gene order: 5’-S (spike) - E (envelope) - M (membrane) - N (nucleocapsid)-3’; numerous small open reading frames, encoding accessory proteins, are distributed among the structural genes (Fig. 1A,B). Some CoVs also encode an additional structural protein, hemagglutinin esterase (HE) (18).

**Figure 1.**
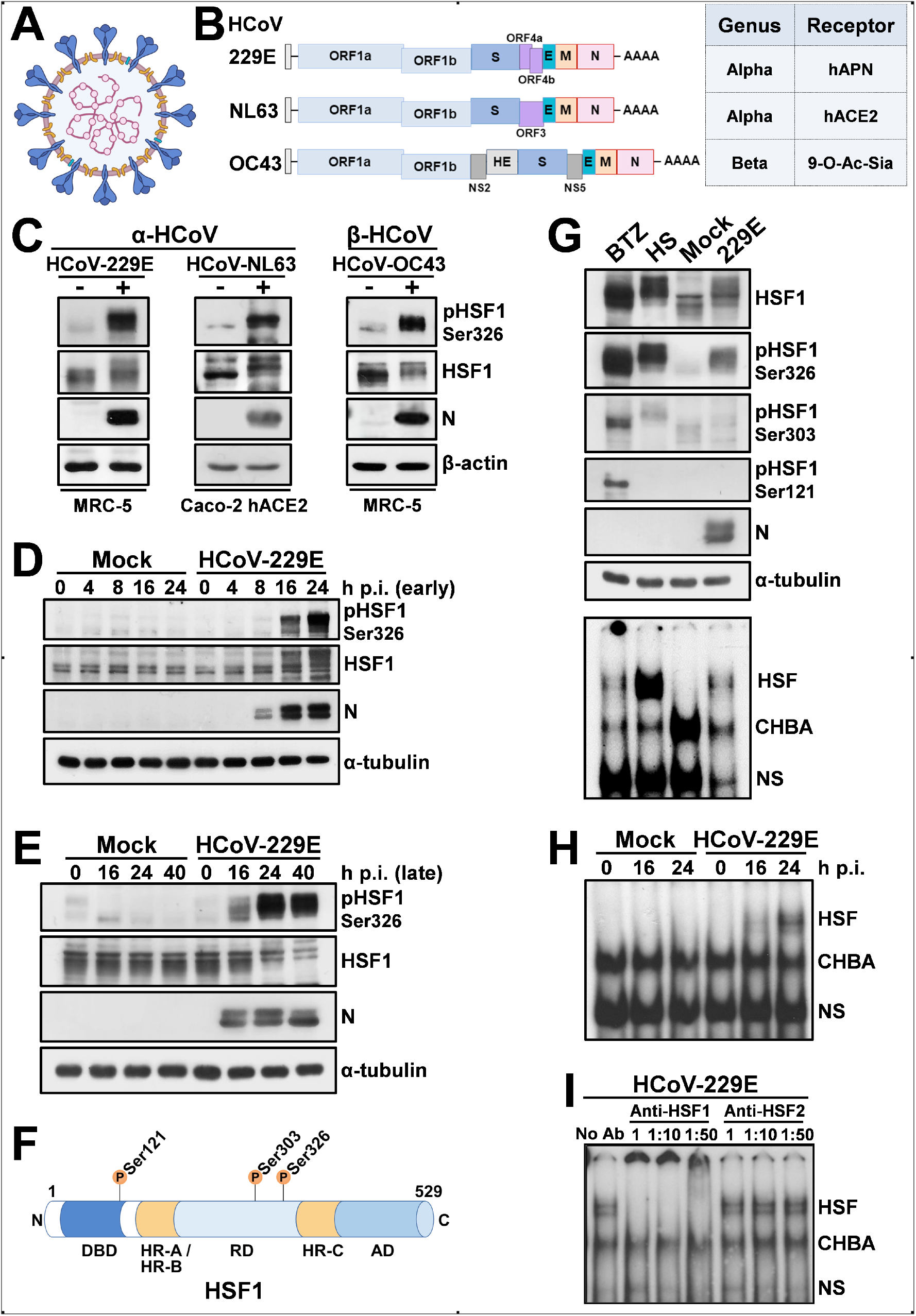
Human coronaviruses induce HSF1 phosphorylation and DNA-binding activity. (**A**) The human coronavirus lipid bilayer comprising the spike protein (blue), the membrane protein (orange) and the envelope protein (green), and the viral RNA (purple) associated with the nucleocapsid protein (pink) are shown. (**B**) Schematic representation of genome structure, classification and receptors of the human coronaviruses HCoV-229E, HCoV-NL63 and HCoV-OC43. ORF1a and ORF1b are represented as light blue boxes; genes encoding structural proteins spike (S), nucleocapsid (N), envelope (E), membrane (M), and hemagglutinin-esterase (HE) and genes encoding accessory proteins are shown. hAPN, human aminopeptidase N; 9-O-Ac-Sia, N-acetyl-9-O-acetylneuraminic acid; ACE2, angiotensin-converting enzyme 2. (**C**) Immunoblot (IB) analysis of pHSF1-Ser326, HSF1, viral nucleocapsid (N) and β-actin protein levels in MRC-5 and Caco-2 hACE2 cells mock-infected (-) or infected (+) with HCoV-229E or HCoV-OC43 (MRC-5) for 24h, or HCoV-NL63 (Caco-2 hACE2) for 72h at a m.o.i. of 0.1 TCID_50_/cell. (**D, E**) Whole-cell extracts (WCE) from samples mock-infected (Mock) or infected with HCoV-229E (1 TCID_50_/cell) were analyzed for pHSF1-Ser326, HSF1, N and α-tubulin protein levels at early (**D**) or late (**E**) times post infection (p.i.) by IB. (**F**) Schematic representation of HSF1 domain organization: DBD, DNA Binding Domain; HR-A/HR-B, Heptad Repeats A and B; RD, Regulatory Domain; HR-C, Heptad Repeat C; AD, Activation Domain. Phosphorylation sites Ser121, Ser303, and Ser326 are shown. (**G**) MRC-5 cells were treated with bortezomib (BTZ, 20 nM) for 16h, exposed to heat stress (HS, 43°C, 40 min), mock-infected (Mock) or infected with HCoV-229E (0.1 TCID_50_/cell) for 40h. WCE were analyzed for levels of HSF1-Ser326, - Ser303 and -Ser121 phosphorylation, HSF1, viral N and α-tubulin proteins by IB (top panels). In the same samples, HSF1 DNA-binding activity was analyzed by EMSA (bottom panel). Positions of the HSF DNA-binding complex (HSF), constitutive HSE-binding activity (CHBA) and nonspecific protein-DNA interaction (NS) are shown. (**H**) MRC-5 cells were mock-infected or infected with HCoV-229E (1 TCID_50_/cell). At different times p.i., HSF1 DNA-binding activity was analyzed by EMSA. (**I**) WCE from samples infected with HCoV-229E (0.1 TCID_50_/cell, 40h p.i.) were preincubated with different dilutions of anti-HSF1 or anti-HSF2 antibodies and analyzed by gel mobility supershift assay. The position of the nonsupershifted virus-induced HSF1 complex is indicated at the left (No Ab).

The genomic RNA complexes with the N protein to form a helical capsid structure surrounded by the viral envelope. Homotrimers of the S protein are embedded in the envelope and extend beyond the viral surface to bind to host receptors giving the virion its crown-like morphology (Fig. 1A). The CoV spike protein is a trimeric class-I fusion glycoprotein (19); each monomer is synthesized as a fusogenically-inactive precursor that assembles into an inactive homotrimer, which is endoproteolytically cleaved by cellular proteases giving rise to a metastable complex of two functional subunits: S1 (bulb) containing the receptor-binding domain responsible for recognition and attachment to the host receptor, and the membrane-anchored S2 (stalk) that contains the fusion machinery (19,20). During synthesis in the infected cell, the nascent spike is imported into the endoplasmic reticulum (ER), where the protein is glycosylated (20,21). S glycoproteins passing the quality control mechanisms of the ER are transported to the ER/Golgi intermediate compartment (ERGIC), the presumed site of viral budding (17–19). M and E proteins are transmembrane proteins, while the HE protein forms smaller spikes in some CoVs (Fig. 1A).

Other typical CoV features include: the expression of many nonstructural genes by ribosomal frameshifting, transcription of downstream genes by synthesis of 3’ nested sub-genomic mRNAs, and the presence of several unusual enzymatic activities encoded within the replicase-transcriptase polyprotein (22). The unique replicative mechanism of CoV involves noncontiguous transcription of the genome, leading to a high rate of recombination, which may play a role in viral evolution and interspecies infections (23).

On the basis of their phylogenetic relationships and genomic structures, CoV are subdivided into four genera: *Alpha-, Beta-, Gamma*- and *Delta-coronavirus*; among these *Alpha*- and *Betα*-CoV infect only mammals, whereas *Gamma*- and *Delta-CoV* infect birds, and only occasionally can infect mammals. Human coronaviruses (HCoV) were discovered in the 1960s and were originally thought to cause only mild disease in humans (17,18). This view changed in 2002 with the SARS (Severe Acute Respiratory Syndrome) epidemic and in 2012 with the MERS (Middle East Respiratory Syndrome) outbreak, two zoonotic infections that resulted in mortality rates greater than 10% and 35%, respectively (17,18). Near the end of 2019, the seventh coronavirus known to infect humans, SARS-CoV-2, phylogenetically in the SARS-CoV clade, emerged in Wuhan, China. SARS-CoV-2 turned out to be a far more serious threat to public health than SARS-CoV and MERS-CoV because of its ability to spread more efficiently, making it difficult to contain worldwide with more than 639 million confirmed cases and over 6.6 million deaths reported worldwide, as of November 30, 2022 (https://covid19.who.int/).

Only two HCoV, HCoV-OC43 and HCoV-229E, were known prior to the emergence of SARS-CoV (24,25), while two more, HCoV-NL63 and HCoV-HKU1, were identified between 2004 and 2005 (26,27). HCoV-OC43 and HCoV-HKU1 likely originated in rodents, while HCoV-229E and HCoV-NL63, similarly to SARS-CoV and MERS-CoV, originated in bats (17). These four HCoV are globally distributed (seasonal CoV, sHCoV) and generally cause only mild upper respiratory diseases (10-30% of all common colds) in immunocompetent hosts, although they can sometimes cause severe and even life-threatening infections especially in infants, elderly people, or immunocompromised patients (28–31). Whereas all seasonal coronaviruses cause respiratory tract infections, HCoV-OC43, HCoV-229E, HCoV-NL63 and HCoV-HKU1 are genetically dissimilar (Fig. 1B), belonging to two distinct taxonomic genera (*Alpha* and *Beta*), and use different receptors that represent the major determinants of tissue tropism and host range (17). HCoV-229E and HCoV-NL63 have adopted cell surface enzymes as receptors, such as aminopeptidase N (APN) for HCoV-229E and angiotensin converting enzyme 2 (ACE2) for HCoV-NL63, while HCoV-OC43 and HCoV-HKU1 use 9-*O*-acetylated sialic acid as a receptor (17,32). In all cases, sHCoV infection is initiated by the binding of the spike (S) glycoprotein, anchored into the viral envelope, to the host receptor (18,22).

At present there is no information on sHCoV infection impact on the host HSR. Herein we show that human coronaviruses trigger a remarkable and sustained activation of HSF1 in late stages of virus replication, leading to the expression of HSF1 canonical and non-canonical target genes in the infected cells. Interestingly, the HSF1-directed transcriptional program turns out to be essential for an efficient virus replication cycle and progeny particles production.

## RESULTS and DISCUSSION

### Human coronaviruses induce HSF1 phosphorylation, nuclear translocation and DNA-binding activity

During a study on the effect of proteostasis disruption in coronavirus-infected cells, we came across the unexpected finding that the human α-CoV 229E was able to trigger the phosphorylation of heat shock factor 1 at serine 326 residue (Ser326), which is considered crucial for HSF1 transcriptional activity (5,33), in human lung cells (Fig. 1C). This accidental finding led us to investigate whether human coronavirus infection results in activation of the HSR in infected cells. We first asked whether HSF1 phosphorylation is only triggered by HCoV-229E or is a common feature of human coronaviruses in different types of cells. Human normal lung MRC-5 fibroblasts and colon carcinoma Caco-2 cells stably expressing the human ACE2 receptor (Caco-2 hACE2) were mock-infected or infected with α-HCoV-229E or β-HCoV-OC43 for 24h and α-HCoV-NL63 (Caco-2 hACE2) for 72h at a m.o.i. (multiplicity of infection) of 0.1 TCID_50_/cell. HCoV-HKU1 was not investigated because of its poor ability to grow in cell culture. At 24 or 72 hours post infection (p.i.), levels of HSF1, the phosphorylated form of HSF1 (pHSF1-Ser326) and the viral nucleocapsid (N) protein as a marker of infection, were determined by immunoblot analysis using specific antibodies in infected cells. As shown in Fig. 1C, all three HCoVs trigger HSF1-Ser326 phosphorylation, as also indicated by slower migration of the factor on SDS polyacrylamide gels.

In order to investigate the dynamic of HSF1 phosphorylation during coronavirus infection, MRC-5 cells were mock-infected or infected with HCoV-229E at a m.o.i. of 0.1 or 1 TCID_50_/cell, and levels of HSF1 phosphorylation were analyzed at different times p.i. Interestingly, high levels of HSF1-Ser326 phosphorylation were detected at late stages of the virus replication cycle, starting at 16h p.i. and continuing up to 40h p.i. (Fig. 1D,E, Supplementary Fig. S1); no Ser326 phosphorylation was detected at early times of infection, up to 12h p.i., independently of the m.o.i. utilized (Fig. 1D, Supplementary Fig. S1B).

According to the PhosphoSitePlus database (https://www.phosphosite.org; October 2021), human HSF1 can be post-translationally modified at 53 residues, including phosphorylation, sumoylation, ubiquitylation and acetylation. Among these, phosphorylation of the HSF1 Regulatory Domain (RD) residues, which is considered a hallmark of HSF1 activity/regulation, turns out to be very complex in human cells as phosphorylation of some sites, in particular Ser326, is pivotal for HSF1 activity, whereas at other sites (e.g., Ser303, in the RD; Ser121 in the DNA-Binding Domain) phosphorylation is associated with attenuation of HSF1 activity (5) (Fig. 1F).

To investigate the outcome of HCoV infection on HSF1 phosphorylation, we compared the effect of two major proteotoxic stressors known to activate HSF1, proteasome inhibition and hyperthermic stress, to HCoV infection on the phosphorylation of different key HSF1 regulatory Ser residues, Ser326, Ser303 and Ser121. MRC-5 cells were treated with the proteasome inhibitor bortezomib (20 nM) for 16h, or exposed to heat stress at 43°C for 40 min, or infected with HCoV-229E (0.1 TCID_50_/cell) for 40h. Whole-cell extracts were analyzed for levels of HSF1, HSF1 phosphorylation at Ser326, Ser121 and Ser303, and viral N protein by immunoblot. In the same samples, HSF1 DNA-binding activity was analyzed by EMSA. As expected, under the conditions utilized, hyperthermic stress resulted in a rapid remarkable increase in HSF1-Ser326 phosphorylation, causing a shift in HSF1 molecular weight, whereas only modestly affected Ser303 and had no effect on Ser121 phosphorylation at this time (Fig. 1G). Phosphorylation of all three serine residues was detected in cells exposed to prolonged (16h) bortezomib treatment, with Ser326 being predominant. On the other hand HCoV-229E infection selectively induced HSF1-Ser326 phosphorylation at levels comparable to heat stress (Fig. 1G). In the same samples, Ser326 phosphorylation was associated with acquisition of HSF1 DNA-binding activity, as shown by EMSA (Fig. 1G, bottom). Concomitantly with HSF1-Ser326 phosphorylation, HSF1 DNA-binding activity was detected at late stages of the virus replication cycle in MRC-5 infected cells, starting at 16h p.i. (Fig. 1H).

Finally, to determine the specificity of HSF1-DNA binding complexes, and whether HCoV also activate HSF2, whole-cell extracts from MRC-5 cells infected with HCoV-229E for 40h were preincubated with different dilutions of anti-HSF1 or anti-HSF2 antibodies and analyzed by gel mobility supershift assay. As in the case of heat shock (12,14), HSF1 was found to be the primary component of the virus-induced HSE-binding activity (Fig. 1I).

As described in the Introduction, in unstressed cells HSF1 is generally found as an inert monomer retained in the cytoplasm in complex with regulatory proteins including HSP90, HSP70, HSP40 and the cytosolic chaperonin complex TRiC (5); upon stress sensing, HSF1 is derepressed in a stepwise process that involves, in addition to post-translational modifications, trimerization and nuclear translocation in order to bind to HSE-containing DNA-sequences (Fig. 2A).

**Figure 2.**
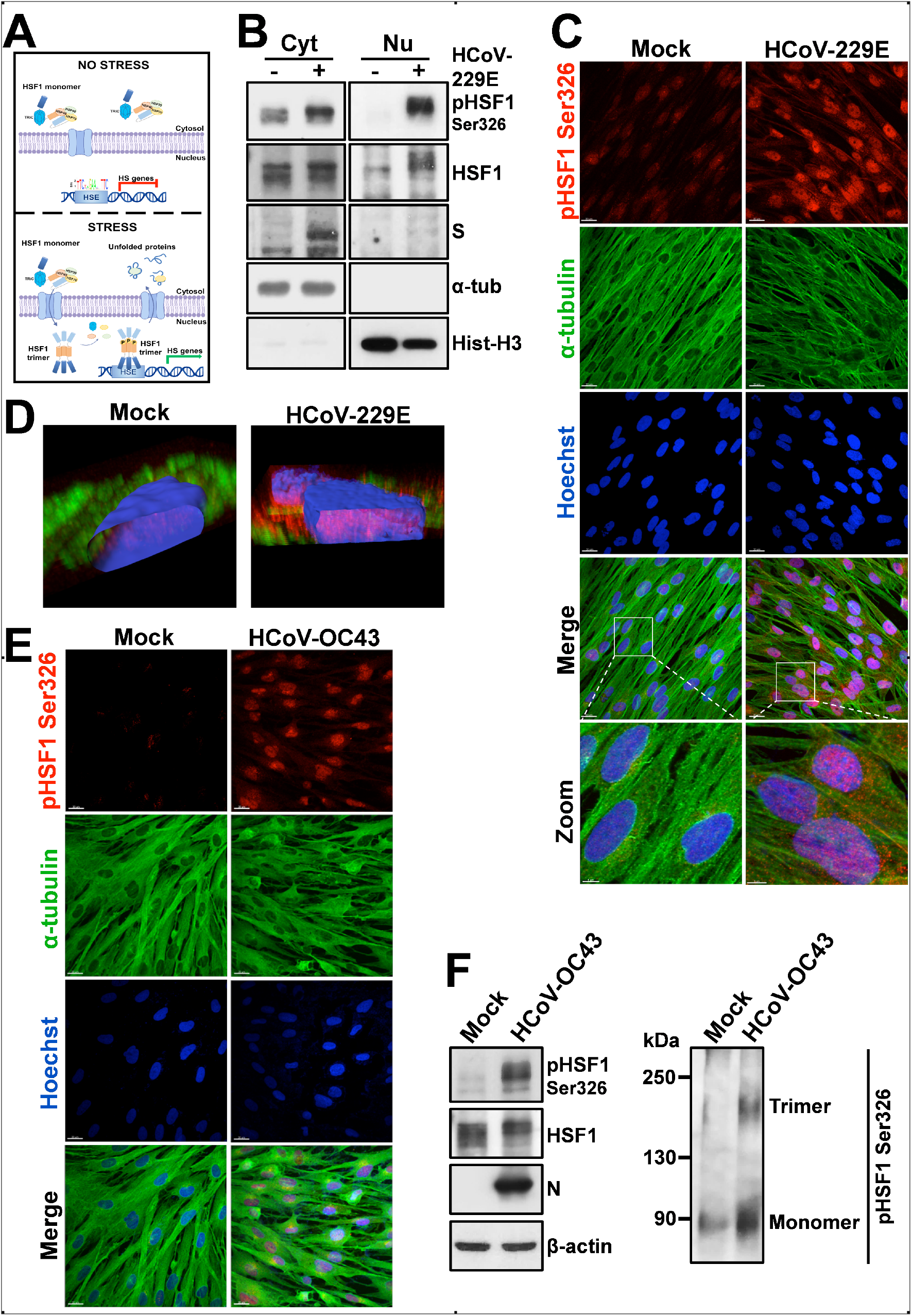
HCoV infection triggers HSF1 nuclear translocation in human lung cells. (**A**) Schematic representation of HSF1 intracellular localization under physiological (no stress) and stress condition. (**B**) Immunoblot analysis of pHSF1-Ser326, HSF1 and viral spike (S) protein levels in cytoplasmic (Cyt) and nuclear (Nu) fractions of MRC-5 cells mock-infected (-) or infected (+) with HCoV-229E (0.1 TCID_50_/cell) for 24h. Antibodies against α-tubulin (α-tub) and histone H3 (Hist-H3) were used as loading controls for cytoplasmic and nuclear fractions, respectively. (**C**) Confocal images of pHSF1-Ser326 (red) and α-tubulin (green) intracellular localization in MRC-5 cells mock-infected or infected with HCoV-229E (1 TCID_50_/cell) at 30h p.i. Nuclei are stained with Hoechst (blue). Merge and zoom images are shown. Scale bar, 20 μm (zoom, 5 μm). (**D**) Confocal 3D-reconstruction of pHSF1-Ser326 (red) intranuclear localization in MRC-5 cells mock-infected or infected as in C; α-tubulin is shown in green. Nuclei are stained with Hoechst (blue). The overlay of the fluorochromes is shown. (**E**) Confocal images of pHSF1-Ser326 (red) and α-tubulin (green) intracellular localization in MRC-5 cells mock-infected or infected with HCoV-OC43 (1 TCID_50_/cell) at 30h p.i. Nuclei are stained with Hoechst (blue). Merge images are shown. Scale bar, 20 μm. (**F**) IB of pHSF1-Ser326, HSF1, N and β-actin protein levels in MRC-5 cells mock-infected or infected with HCoV-OC43 (0.1 TCID50/cell) for 24h (left panels). HSF1 monomers and trimers in the same samples are shown (right panel).

To investigate whether HCoV infection causes intracellular redistribution of the factor, HSF1 localization was determined by cell-fractionation and confocal-immunomicroscopy studies in MRC-5 lung cells infected with the α-HCoV-229E (0.1 TCID_50_/cell) for 24h. As shown in Fig. 2B,C, under normal conditions HSF1 is predominantly localized in the cytoplasm of MRC-5 cells; HCoV-229E infection, in addition to Ser326 phosphorylation, caused a dramatic redistribution of the factor, which is found in the nuclei of infected cells in a remarkably abundant amount (Fig. 2C,D). Similar results were obtained in MRC-5 cells infected with the β-HCoV-OC43 that caused, in addition to Ser326 phosphorylation, HSF1 trimerization and translocation to the nucleus (Fig. 2E,F). These results demonstrate that both *Alpha*- and *Beta*-coronaviruses trigger HSF1 phosphorylation and nuclear translocation.

### HCoV infection turns on an HSF1-driven transcriptional program in human lung cells

Since phosphorylation, nuclear translocation and DNA-binding activity in some instances may not warrant target genes transcription (34,35), we next investigated whether HCoV-induced nuclear HSF1 is transcriptionally active. HSF1-target gene expression was compared in MRC-5 cells mock-infected or infected with HCoV-229E (0.1 TCID50/cell) for 24h using a qRT-PCR array (PAHS-076ZD-2 – Qiagen), which profiles the expression of 84 HS genes encoding HSPs and molecular chaperones. As shown in Fig. 3A,B, HCoV infection resulted in the expression of high levels of several HS genes, demonstrating that the virus-induced HSF1 is transcriptionally active.

**Figure 3.**
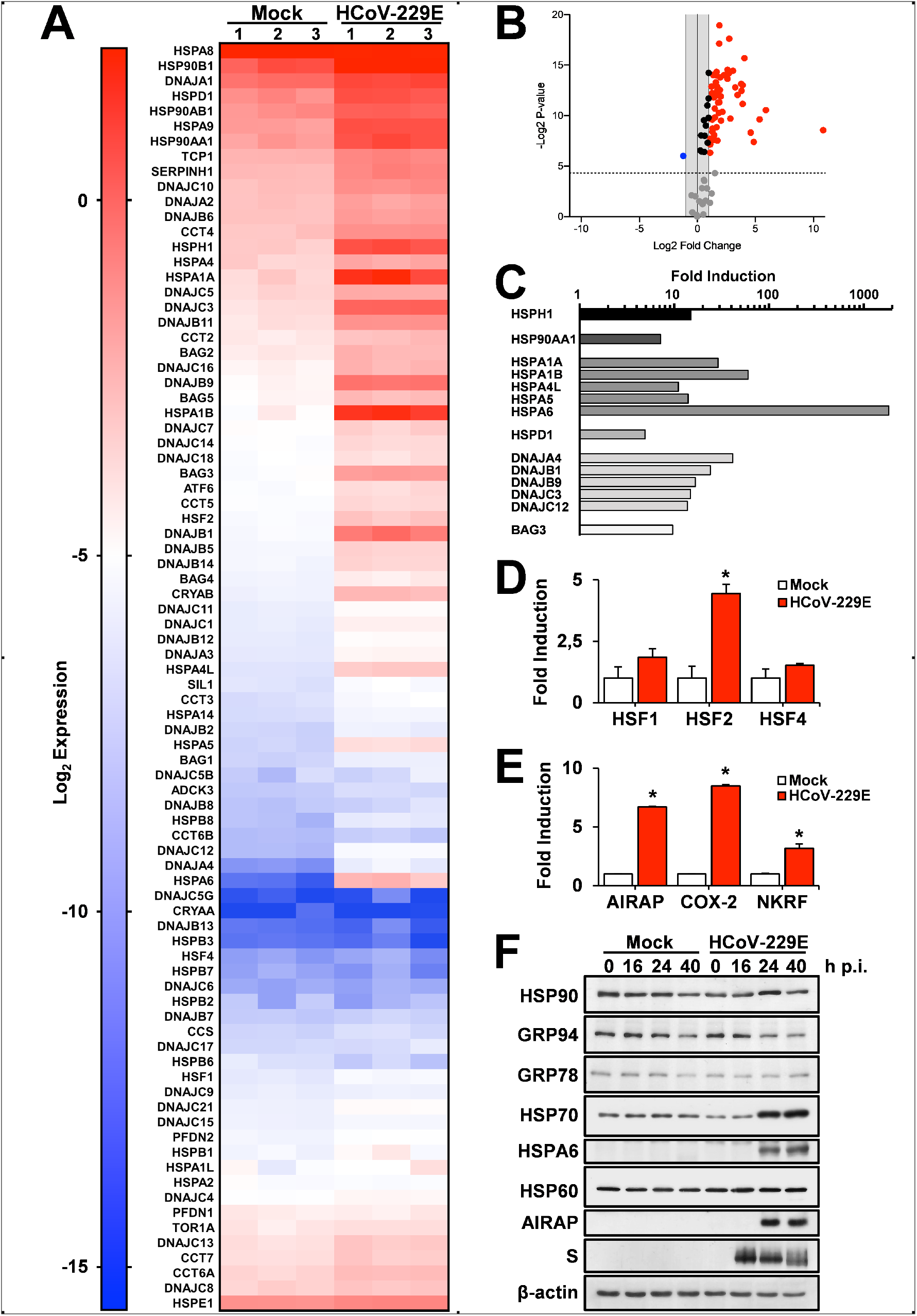
HCoV infection turns on an HSF1-driven transcriptional program in human lung cells. (**A-D**) Expression profile of selected HSF1-target genes affected by HCoV-229E infection (0.1 TCID50/cell) for 24h in MRC-5 cells relative to mock-infected cells as determined by microarray analysis. Heat Map (A) and Volcano plot (B) of 84 human heat shock protein and chaperones/co-chaperones gene expression. In (A) each row represents a single gene, each column represents a sample [mock-infected (Mock) or HCoV-229E infected cells; n=3]. The gradual color ranging from blue to red represents the mRNA expression level. In the Volcano plot (B) fold regulation threshold is set to 2 and *p*-value cut off is 0.05; each dot represents a gene: red dots indicate significantly upregulated genes and blue dots indicate significantly downregulated genes. Selected heat shock proteins and chaperones/co-chaperones genes whose expression is highly induced by HCoV infection are shown in (C); levels of heat shock factors (HSF1, HSF2 and HSF4) gene expression affected by HCoV infection are shown in (D). (**E**) Expression of non-canonical HSF1-target genes AIRAP, COX-2 and NKRF in samples treated as in A as determined by qRT-PCR. Error bars indicate means± S.D. * = p < 0.05; Student’s *t*-test (D, E). (**F**) Levels of HSP90, GRP94, GRP78, HSP70, HSPA6, HSP60, AIRAP, viral spike (S) and β-actin proteins were determined by immunoblot in MRC-5 cells mock-infected or infected with HCoV-229E (1 TCID_50_/cell) at different times p.i.

Whereas HSF1 has been generally considered to function in the classical heat shock response as a guardian of cellular health, accumulating evidence have challenged these traditional views, and HSF1 has been shown to drive diverse transcriptional programs in development, metabolism and cancer that are distinct from the classical HSR (36–38). In the case of the HCoV-driven response we found that viral infection resulted in the expression of high levels of different HSP mRNAs belonging to the HSP70 and HSP90 families of chaperones, as well as glucose-regulated proteins GRP78/BiP (HSPA5), GRP94 (HSP90B1) and a plethora of other HSF1-target genes. The expression of molecular chaperones of the DNAJ (HSP40) family, characterized by a highly conserved amino acid stretch known as the ‘J-domain’ and with critical functions in protein folding and oligomeric protein complex assembly, is also notably increased; in particular, DNAJA4 expression was increased more than 40-fold, and DNAJB1 (Hdj1/Sis1), DNAJB9 (ERdj4), DNAJC3 (ERdj6) and DNAJC12 more than 10-fold (Fig. 3C). The expression of the HSPH1 gene (member of the HSP110 family) was also elevated, whereas genes belonging to the HSP60 (HSPD1) family were increased to a lesser extent (Fig. 3A,C). Extremely high levels were detected for the HSPA6 gene product, whose expression was increased more than 1800-fold; notably, HCoV infection also strongly (>9-fold) increased the expression of the HSP70 cochaperone BAG3 (Bcl-2-associated athanogene domain 3), a multidomain protein with anti-apoptotic function known to play a central role in autophagy and involved in the regulation of the dynein motor pathway (Fig. 3C) (39,40).

Furthermore, whereas HCoV infection did not affect HSF1 or HSF4 expression, it resulted in a significant increase of HSF2 expression (Fig. 3A,D); these results confirm the recent observation that HSF1 promotes HSF2 gene transcription in human cells, representing an interesting example of transcription factor involved in controlling the expression of members of the same family (41).

As indicated above, beyond HS genes, HSF1-binding sites have been described in several genes encoding proteins with non-chaperone function (38,42). We have previously identified three non-canonical HSF1-target genes, characterized by the presence of HSE elements in the promoter of the human (but not the murine) gene: 1) the zinc-finger AN1-type domain-2a gene, also known as AIRAP (arsenite-inducible RNA-associated protein) (8,14), which was recently identified as a regulator of prosurvival networks in melanoma cells, interacting and stabilizing the anti-apoptotic protein cIAP2 (43); 2) NKRF (NF-κB repressing factor), known as a silencer of NF-κB, a transcription factor with a critical role in the control of inflammation, cell survival and virus replication (44,45), which was recently identified as an unconventional nucleolar HSP crucial for correct ribosomal RNA processing and for preventing aberrant rRNA precursors and discarded fragment accumulation (15); 3) human cyclooxygenase-2 (hCOX-2), a key regulator of inflammation controlling prostanoid and thromboxane synthesis, which was found to be temperature-regulated in human primary cells via a distal *cis*-acting HSE located at position −2495 from the transcription start site (12).

We therefore investigated whether HCoV infection induced the expression of AIRAP, NKRF and COX-2 by qRT-PCR analysis. As shown in Fig. 3E, the expression of all three genes was found to be significantly increased in MRC-5 cells at 24h after infection with HCoV-229E (0.1 TCID_50_/cell).

Notably, the expression of HSF1-target genes, including HSP70 and AIRAP, was dependent on the virus m.o.i. with levels higher than 1000-fold for HSP70 when cells were infected at a m.o.i. of 1 TCID_50_/cell for 24h (Supplementary Fig. S2A). Furthermore, very high levels of HSF1-target genes expression were detected up to 40h after infection (Supplementary Fig. S2B).

Similarly to many RNA viruses, CoV infection is known to affect the host cell translational machinery and turn off cellular protein synthesis to promote viral RNA translation employing, in addition to cap-dependent translation, non-canonical translation mechanisms to expand their coding capacity such as ribosomal frameshifting and ribosomal shunting (22,46); therefore, induction of the expression of cellular genes, even at high levels, not necessarily may lead to an increase in the relative protein levels. We then investigated whether HCoV-induced HSF1 activation resulted in the expression of a set of canonical and non-canonical HSP. The results, shown in Fig. 3F, indicate that the level of some, but not all, the proteins examined was increased in infected cells. In particular, levels of HSP70, HSPA6 and AIRAP were remarkably high in HCoV-229E-infected MRC-5 cells at late stages of viral replication, whereas no significant changes in the levels of HSP90, HSP60, GRP78 and GRP94 were detected at all times examined.

Finally, similarly to HCoV-229E, infection with both HCoV-OC43 and HCoV-NL63 triggered the expression of high levels of HSF1-dependent proteins, independently of the host cell type (Supplementary Fig. S3).

These results demonstrate that HCoV trigger a powerful and distinct HSF1-driven transcriptional/translational response in infected cells at late stages of infection and prompted us to investigate whether this phenomenon only reflects a cellular defense response to the invading pathogen, or whether the virus activates and hijacks the HSF1-pathway for its own gain.

### HSF1 activation is required for efficient replication of human coronaviruses

We first investigated the effect of HSF1-silencing on HCoV-229E infection. In a first set of experiments MRC-5 cells were transiently transfected with two different HSF1 siRNA (siHSF1_1_ and siHSF1_2_) or scramble-RNA. After 48h, cells were infected with HCoV-229E (0.1 TCID_50_/cell) and, at 24h p.i., whole-cell extracts were analyzed for levels of HSF1, pHSF1-Ser326, the HSF1 target gene AIRAP and the viral spike (S) protein by immunoblot, while virus yield was determined in cell supernatants. An efficient HSF1-silencing was obtained with both HSF1-siRNAs, as confirmed also by the lower levels of AIRAP in HSF1-silenced infected cells (Fig. 4A, top). Interestingly, HSF1-silencing also resulted in decreasing viral spike levels (Fig. 4A, top); concomitantly, infectious viral progeny production was significantly reduced in the supernatants of infected cells after silencing with both HSF1-siRNAs (Fig. 4A, bottom), suggesting that HSF1 may be necessary for optimal virus replication. Next, we analyzed HCoV replication in stably HSF1-silenced HeLa cells (HeLa-HSF1i) as compared to wild-type (HeLa-WT) cells (47). Because of the lack of appropriate receptors for α- and β-CoVs in these cells, 229E and OC43 HCoV genomic RNA was extracted from infectious virions and transfected into HeLa-HSF1i or HeLa-WT monolayers together with a GFP reporter plasmid as internal transfection control for 4h. After 48h or 72h, levels of HSF1, GFP, and viral S and N proteins were analyzed in cell extracts by immunoblot, while virus yield was determined in the supernatant of transfected cells. As shown in Fig. 4B and Supplementary Fig. S4A,B, genomic RNA transfection resulted in the production of infectious viral progeny in HeLa cells at 48h or 72h after transfection. Notably, both HCoV-229E and HCoV-OC43 virus yield and viral structural protein levels were significantly lower in HeLa HSF1i cells as compared to wild-type cells (Fig. 4B and Supplementary Fig. S4A,B), confirming that HSF1 is needed for both α- and β-HCoV efficient replication and indicating that HSF1 inhibitors may counteract human coronavirus infection.

**Figure 4.**
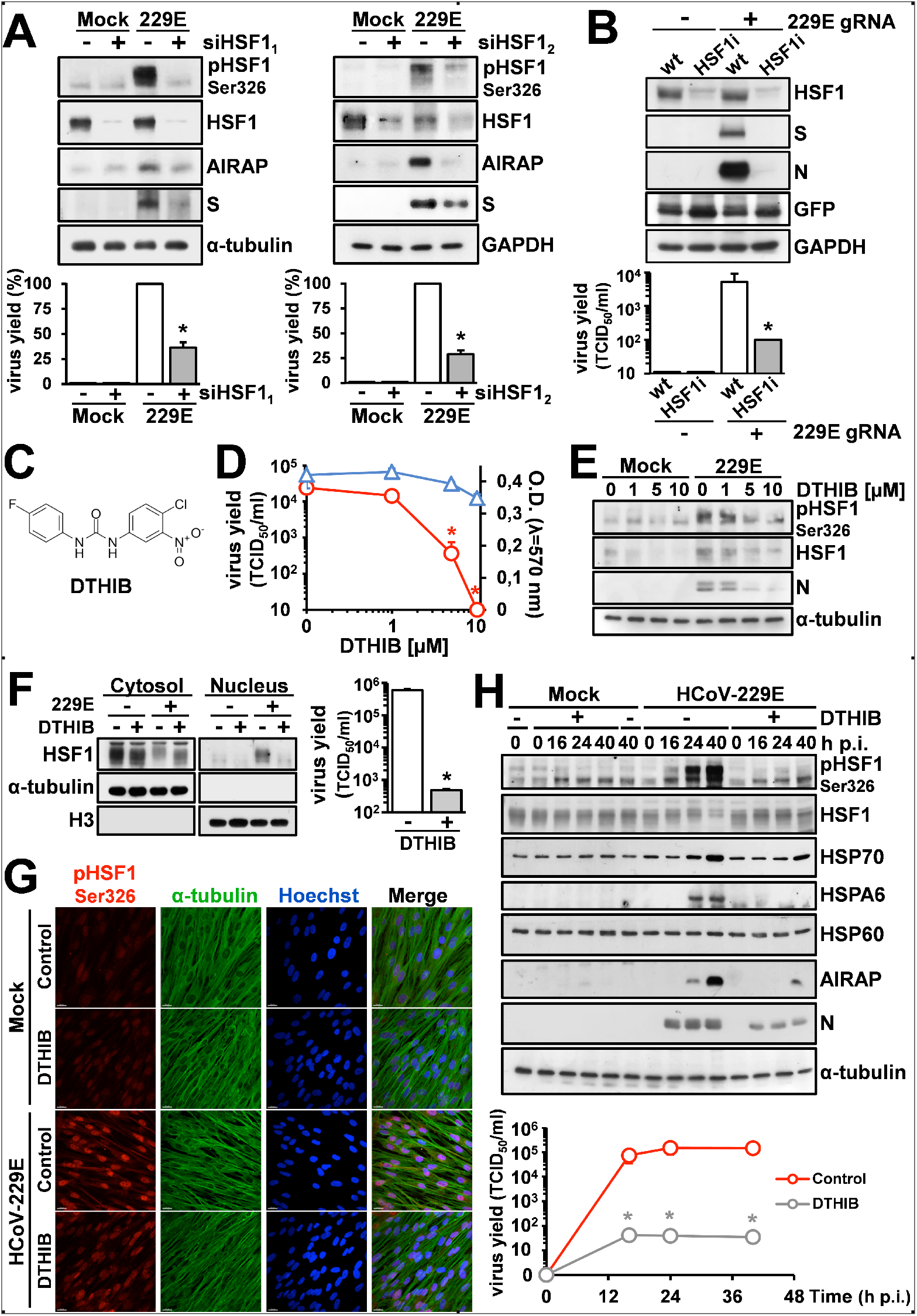
HSF1 activation is required for efficient replication of human coronaviruses. **(A)** Immunoblot of pHSF1-Ser326, HSF1, AIRAP, viral spike (S), α-tubulin and GAPDH protein levels in MRC-5 cells transiently transfected with two different HSF1-siRNAs [siHSF1_1_ (left) and siHSF1_2_ (right); +] or scramble-RNA (-) for 48h, and infected with HCoV-229E (0.1 TCID_50_/cell) or mock-infected (Mock) for 24h (top panels). Virus yield was determined at 24h p.i. by TCID_50_ infectivity assay (bottom panel). Data, expressed as percentage of control, represent the mean ± S.D. of duplicate samples. * = p < 0.05; Student’s *t*-test. (**B**) Wild type (wt) or stably HSF1-silenced (HSF1i) HeLa cells were co-transfected with HCoV-229E genomic RNA (229E gRNA) and the pCMV-GFP vector for 4h. After 48h, levels of HSF1, viral spike (S) and nucleocapsid (N), GFP and GAPDH proteins were analyzed by IB (top panels). Virus yield in the supernatant of transfected cells was determined by TCID50 infectivity assay (bottom panel). Data, expressed as TCID50/ml, represent the mean ± S.D. of duplicate samples. * = p < 0.05; Student’s *t*-test. (**C**) Structure of DTHIB (Direct Targeted HSF1 Inhibitor). (**D**) MRC-5 cells mock-infected or infected with HCoV-229E (0.1 TCID_50_/cell) were treated with different concentrations of DTHIB immediately after the adsorption period. Virus yield (**O**) was determined at 24h p.i. by TCID_50_ infectivity assay. Data, expressed as TCID_50_/ml, represent the mean ± S.D. of duplicate samples. * = p < 0.05; ANOVA test. In parallel, cell viability (Δ) was determined in mock-infected cells by MTT assay. Absorbance (O.D.) of converted dye was measured at λ = 570nm. (E) Immunoblot of pHSF1-Ser326, HSF1, viral N and α-tubulin protein levels in samples treated as in D. (F) Western blot analysis of HSF1 protein levels in cytoplasmic (Cytosol) and nuclear (Nucleus) fractions of MRC-5 cells mock-infected (-) or infected (+) with HCoV-229E (0.1 TCID50/cell) for 24h and treated with DTHIB (10 μM, +) or vehicle (-) immediately after the adsorption period (left panel). Antibodies against α-tubulin and Histone H3 (H3) were used as a loading control for cytoplasmic and nuclear fractions, respectively. Virus yield was determined at 24h p.i. by TCID_50_ infectivity assay (right panel). Data, expressed as TCID_50_/ml, represent the mean ± S.D. of duplicate samples. * = p < 0.05; Student’s *t*-test. (G) Confocal images of pHSF1-Ser326 (red) and α-tubulin (green) intracellular localization in MRC-5 cells mock-infected or infected with HCoV-229E (1 TCID50/cell) for 30h and treated with DTHIB (5 μM) or control diluent immediately after the adsorption period. Nuclei are stained with Hoechst (blue). Merge images are shown. Scale bar, 20 μm. (H) MRC-5 cells were mock-infected or infected with HCoV-229E (1 TCID50/cell) and treated with DTHIB (7,5 μM). At different times p.i., levels of pHSF1-Ser326, HSF1, HSP70, HSPA6, HSP60, AIRAP, viral N and α-tubulin proteins were determined by IB (top panels). In parallel, virus yield was determined by TCID50 infectivity assay (bottom panel). Data, expressed as TCID_50_/ml, represent the mean ± S.D. of duplicate samples. * = p < 0.05; Student’s *t*-test.

Different inhibitors of the HSF1 pathway have been recently described (48). An interesting new approach to selectively target nuclear HSF1 without disrupting the HSR cytoplasmic signaling, is represented by DTHIB (Direct Targeted HSF1 InhiBitor) (Fig. 4C), which physically engages the HSF1 DNA binding domain (DBD) and has been shown to selectively stimulate degradation of the factor in the nucleus of prostate cancer cells (49).

In order to ascertain whether DTHIB could downregulate HCoV-induced nuclear HSF1 in lung cells and determine its effect on HCoV replication, MRC-5 cells mock-infected or infected with HCoV-229E (0.1 TCID_50_/cell) were treated with different concentrations of DTHIB immediately after the virus adsorption period, and virus yield was determined at 24h p.i. by infectivity assay. In parallel, the effect of the drug on mock-infected MRC-5 cell viability was determined by MTT assay. As shown in Fig. 4D, DTHIB was found to be very effective in inhibiting HCoV-229E replication causing a >3-log reduction in infectious viral particle production at non cytotoxic concentrations. This effect was accompanied by a decrease in the level of HSF1-Ser326 phosphorylation and viral N protein expression (Fig. 4E), as well as nuclear (but not cytoplasmic) levels of HSF1 (Fig. 4F,G), as previously reported (49). Similar results were obtained in human cells infected with HCoV-OC43 (Supplementary Fig. 4C), demonstrating that DTHIB is effective in inhibiting both α- and β-HCoV replication.

Finally, to investigate the effect of DTHIB on the dynamic of HCoV-induced HSR activation, HCoV-infected MRC-5 cells were treated with DTHIB (7,5 μM) and levels of pHSF1-Ser326, HSF1, HSP70, HSPA6, HSP60, AIRAP, viral N and α-tubulin proteins were analyzed at different times p.i. Results, shown in Fig. 4H, confirm that DTHIB treatment inhibits virus-induced HSF1-dependent gene expression and viral replication up to 40h p.i.

## CONCLUSIONS

As described in the Introduction the four known seasonal coronaviruses HCoV-229E, HCoV-NL63, HCoV-OC43 and HCoV-HKU1, are globally circulating in the human population and are characterized by their inability to induce long-lasting protective immunity (50). Although sHCoV generally cause only mild upper respiratory diseases in immunocompetent hosts, they can sometimes cause severe lower respiratory infections, including life-threatening pneumonia and bronchiolitis (28–31). Besides respiratory illnesses, sHCoV may cause enteric and neurological diseases (51–54), while a possible involvement of HCoV-229E in the development of Kawasaki disease was suggested (55,56); in addition, HCoV-OC43 has been shown to have neuroinvasive properties and to cause encephalitis in animal models (57,58). There is no specific treatment for sHCoV infections, also due to the limited knowledge on sHCoV biology.

As indicated above, coronavirus infection is initiated by the binding of the spike glycoprotein to the host receptor (17,22). After virus entry into cells, the positive-sense RNA genome hijacks the host ribosomes acting as a direct template for protein translation of two large open reading frames, ORF1a and ORF1b, into the polyproteins pp1a and pp1ab. Pp1a and pp1ab are co-translationally and post-translationally processed by the ORF1a-encoded proteases into different non-structural proteins (nsps) that form the multi-subunit RNA replicase–transcriptase complex (RTC) containing several nsps, including the RNA-dependent RNA polymerase (RdRp), the helicase and the exonucleaseN proteins (17,22). The RTC machinery needs to localize to modified intracellular membranes derived from the ER in the perinuclear region, where it drives the generation of negative-sense RNAs [(–)RNAs]. During replication, full-length (–)RNA copies of the genome are synthesized and used as templates for the production of progeny RNA genomes; during transcription, a subset of 7–9 subgenomic RNAs, including those encoding structural proteins, is produced through discontinuous transcription, followed by synthesis of viral mRNAs and translation of viral proteins. A quite complex and yet not well deciphered mechanism of assembly of both ribonucleocapsid structures and viral envelopes requires intense intracellular trafficking to and from the ER and the ERGIC, followed by release of the newly produced coronavirus particle from the infected cell. Understanding the host-virus interactions enabling cells to sustain all the complex steps of the coronavirus lifecycle is pivotal to identifying potential targets for host-directed antiviral strategies.

We now show that both *Alpha*- and *Beta*-human coronaviruses are potent inducers of the proteostasis guardian HSF1, by selectively promoting HSF1 phosphorylation at serine 326 residue, which is crucial for HSF1 transcriptional activity, and leading to HSF1 nuclear translocation, binding to HSE sequences and triggering a powerful and distinct HSF1-driven transcriptional/translational response in infected cells at late stages of infection. Despite the fact that, as indicated above, coronavirus infection is known to affect the host cell translational machinery and turn off cellular protein synthesis to promote viral RNA translation (22,46), remarkably high levels of some, but not all, canonical and non-canonical HSF1-target genes products, including HSP70, HSPA6 and AIRAP, were found in HCoV-infected cells at late stages of the viral lifecycle.

Interestingly, silencing experiments demonstrate that activation of the HSR does not merely reflect a cellular response to the virus-induced proteotoxic stress caused by the abundant synthesis and intracellular trafficking of foreign proteins, but that HCoV activate and hijack the HSF1-pathway for their own gain. Notably, post infection treatment with the recently described HSF1 inhibitor DTHIB, that selectively targets nuclear HSF1 physically engaging the factor DNA-binding domain and stimulating its degradation, was found to be highly effective in inhibiting sHCoV replication, remarkably reducing the production of HCoV infectious progeny particles in lung cells. Whereas the role of different canonical and non-canonical HSPs in HCoV replication remains to be elucidated, these results open new scenarios for the search of innovative antiviral strategies in the treatment of coronavirus infections.

## MATERIALS and METHODS

### Cell culture and treatments

Human normal lung MRC-5 fibroblasts (American Type Culture Collection, ATCC, CCL-171), Caco-2 hACE2 cells, stably expressing the hACE2 receptor, and HeLa cells, stably transfected with pSUPER-HSF1i/pcDNA (HeLa-HSF1i) or control (HeLa wild-type) plasmids, were grown at 37°C in a 5% CO_2_ atmosphere in EMEM (MRC-5 cells) or DMEM (Caco-2 hACE2 and HeLa cells) medium, supplemented with 10% fetal calf serum (FCS), 2 mM glutamine and antibiotics. Generation of Caco-2 hACE2 cells (59) and HeLa-HSF1i cells (47) was described previously. For heating procedures, cells were subjected to heat shock at the indicated temperature in a precision water bath W14 (Grant Instruments) (15). DTHIB (Direct Targeted HSF1 Inhibitor) (Selleckchem) dissolved in DMSO stock solution, was diluted in culture medium, added to infected cells after the virus adsorption period, and maintained in the medium for the duration of the experiment. Bortezomib (BTZ, Selleckchem) was dissolved in DMSO and diluted in culture medium immediately before use. Controls received equal amounts of DMSO vehicle, which did not affect cell viability or virus replication. Cell viability was determined by the 3-(4,5-dimethylthiazol-2-yl)-2,5-diphenyltetrazolium bromide (MTT) to MTT formazan conversion assay (Sigma-Aldrich), as described (60). The 50% lethal dose (LD50) was calculated using Prism 5.0 software (Graph-Pad Software Inc.). Microscopical examination of mock-infected or virus-infected cells was performed daily to detect virus-induced cytopathic effect and possible morphological changes and/or cytoprotection induced by the drug. Microscopy studies were performed using a Leica DM-IL microscope and images were captured on a Leica DC 300 camera using Leica Image-Manager500 software.

### Coronavirus infection and titration

Human coronaviruses HCoV-229E (ATCC), HCoV-OC43 (ATCC) and HCoV-NL63 (strain Amsterdam-1, a kind gift from Lia van der Hoek, University of Amsterdam), were used for this study (61). For virus infection, confluent MRC-5 (229E and OC43) or Caco-2 hACE2 (NL63) cell monolayers were infected with HCoV for 1 (229E and OC43) or 2 (NL63) hours at 33°C at a multiplicity of infection (m.o.i.) of 0.1 or 1 TCID_50_ (50% tissue culture infectious dose)/cell. After the adsorption period, the viral inoculum was removed, and cell monolayers were washed three times with phosphate-buffered saline (PBS). Cells were maintained at 33°C in growth medium containing 2% FCS. Virus yield was determined at different times after infection by TCID50 infectivity assay, as described previously (62). The 50% inhibitory concentration (IC_50_) of the compound tested was calculated using Prism 5.0 software.

### Protein Analysis and Western blot

For analysis of proteins, whole-cell extracts (WCE) were prepared after lysis in High Salt Buffer (HSB) (59). Nuclear and cytoplasmic extracts were prepared as described (63). For Western blot analysis, cell extracts (15 μg/sample) were separated by SDS-PAGE under reducing or non-reducing conditions and blotted to a nitrocellulose membrane. Membranes were incubated with the selected antibodies, followed by incubation with peroxidase-labeled anti-rabbit, anti-mouse or anti-goat IgG. Primary and secondary antibodies used are listed in Supplementary Table S1. Quantitative evaluation of proteins was determined as described (64).

### Electrophoretic mobility shift assay (EMSA)

The 35-bp HSP70-HSE DNA probe was described previously (65). WCE (10 μg) prepared after lysis in high-salt extraction buffer were incubated with a ^32^P-labeled HSE DNA probe (66) followed by analysis of DNA-binding activity by EMSA. Binding reactions were performed as described (67). Complexes were analyzed by nondenaturing 4% polyacrylamide gel electrophoresis. To determine the specificity of HSF-DNA complexes, WCE were preincubated with different dilutions of anti-HSF1 (sc-9144, Santa Cruz Biotechnology) or anti-HSF2 (sc-13056, Santa Cruz Biotechnology) antibodies for 15 min before electromobility supershift assay (68).

### Immunofluorescence microscopy

MRC-5 cells infected with HCoV-OC43 or HCoV-229E were grown in 8-well chamber slides (Lab-Tek II) and, after the adsorption period, were treated with DTHIB, or vehicle for 30h. Cells were fixed, permeabilized and processed for immunofluorescence as described (59) using selected antibodies, followed by decoration with Alexa Fluor 555- or 488-conjugated antibodies (Molecular Probes, Invitrogen). Nuclei were stained with Hoechst 33342 (Molecular Probes, Invitrogen). For confocal microscopy, images were acquired on Olympus FluoView FV-1000 confocal laser scanning system (Olympus America Inc., Center Valley, PA) and analyzed using Imaris (v6.2) software (Bitplane, Zurich, Switzerland). Images shown in all figures are representative of at least three random fields (scale-bars are indicated).

### RNA extraction, gene expression and Real-time PCR analysis

Total RNA from mock-infected or infected cells was prepared using ReliaPrep RNA Cell Miniprep System (Promega) and reverse transcription was performed with PrimeScript RT Reagent Kit (Takara) according to the manufacturer’s protocol. Gene expression of 84 heat shock genes in mock-infected or 229E-infected cells was analyzed using Human Heat Shock Proteins & Chaperones RT^2^ Profiler PCR array (Qiagen). Real-time PCR analyses were performed with specific primers, listed in Supplementary Table S2, using SsoAdvanced Universal SYBR Green Supermix (CFX96, Bio-Rad). Relative quantities of selected mRNAs were normalized to L34 (61). All reactions were made in triplicate using samples derived from at least three biological repeats.

### siRNA interference

Two siRNA duplex target sequences (siHSF1_1_ and siHSF1_2_) and their scrambled control (scrRNA) (QIAGEN) were used for HSF1-silencing (Supplementary Table S2). Transfections were performed using jetPRIME Transfection Reagent, according to the manufacturer’s instructions. In brief, cells were plated on 35-mm wells (2,5 × 10^5^ cells/well) and, after 24h, were transfected with 50 nM of the indicated siRNAs (siHSF1_1_ and siHSF1_2_) or scrRNA. After 24h, cells were washed twice with culture medium and transfection was repeated as above. At 24h after the second transfection, siRNAs were removed, and cells were washed twice with culture medium before HCoV-229E infection.

### HCoV genomic RNA transfection

For HCoV genomic RNA transfection experiments, MRC-5 cell monolayers were infected with HCoV-229E or HCoV-OC43 for 1h at 33°C at an m.o.i. of 0.1 TCID_50_/cell and HCoV genomic RNA was extracted from the supernatants at 24h p.i. using TRIzol-LS reagent (Life Technologies) as described in the manufacturer protocol. HeLa wild-type or HeLa-HSF1i cell monolayers were mock-transfected or co-transfected with HCoV genomic RNA (1 μg/ml) and pCMV-GFP vector (Clontech) using TransIT-mRNA Transfection Kit (Mirus Bio) at 33°C (61). After 4h, transfection medium was removed and cells were maintained at 33°C in growth medium containing 2% FCS. After 48h (229E) or 72h (OC43), culture supernatants were collected for virus progeny titer determination, and cell monolayers were processed by Western blot analysis.

### Statistical analysis

Statistical analyses were performed using Prism 5.0 software (GraphPad Software). Comparisons between two groups were made using Student’s *t*-test; comparisons among groups were performed by one-way ANOVA with Bonferroni adjustments. *p* values ≤0.05 were considered significant.

Data are expressed as the means ± standard deviations (S.D.) of results from duplicate or quadruplicate samples. Each experiment (in duplicate) was repeated at least twice.

## Author Contributions

S. Pauciullo, A. Riccio, A. Rossi, SS and S. Piacentini performed the experiments; MGS designed the study; MGS, S. Pauciullo and A. Riccio wrote the manuscript. All authors contributed to the interpretation of the data and approve the content of the manuscript.

## Acknowledgments

The authors thank Lia van der Hoek (Academic Medical Center, University of Amsterdam) for providing the HCoV-NL63 Amsterdam-1 strain, and Dennis Thiele (Department of Pharmacology and Cancer Biology, Duke University Durham, NC) for the useful discussion. Silvia Pauciullo is enrolled in the PhD Program in Cellular and Molecular Biology, Department of Biology, University of Rome Tor Vergata, Rome, Italy.

## Data availability statement

Data will be made available on reasonable request

