## Supplemental Figures for "Human coronaviruses activate and hijack the proteostasis guardian HSF1 to enhance viral replication"

### Supplementary Table S1. Antibodies used

| Antibody | Source | Catalogue Number |
|---|---|---|
| HCoV-229E Spike (P) | LGC NAC Company | PAB21477-100 |
| HCoV-229E Nucleocapsid (P) | Sino Biological | 40640-T62 |
| HCoV-OC43 Nucleocapsid (P) | Sino Biological | 40643-T62 |
| KDEL (M) | Enzo Life Sciences | ADI-SPA-827 |
| HCoV-NL63 Nucleocapsid (P) | Sino Biological | 40641-T62 |
| α-Tubulin (M) | Sigma-Aldrich | T5168 |
| HSF1 (P) | Enzo Life Sciences | ADI-SPA-901 |
| HSF1 (phospho-S121) (P) | Tebubio | A8041 |
| HSF1 (phospho-S303) (P) | abcam | ab47369 |
| HSF1 (phospho-S326) (P) | abcam | ab76076 |
| β-Actin (P) | Sigma-Aldrich | A2066 |
| ZFAND2A (P) | Sigma-Aldrich | HPA019469 |
| GAPDH (P) | Cusabio | CSB-PA00025A0Rb |
| HSP70/HSP72 (M) | Enzo Life Sciences | ADI-SPA-810 |
| Histone H3 (P) | abcam | ab1791 |
| HSP90β (P) | Santa Cruz Biotechnology | sc-1057 |
| HSPA6 (P) | Enzo Life Sciences | ADI-SPA-756 |
| HSP60 (P) | StressMarq | SPC-105 |
| GFP (M) | CUSABIO | CSB-MA000051M0m |
| Alexa Fluor 488 goat anti-mouse | Invitrogen | A11001 |
| Alexa Fluor 555 goat anti-rabbit | Invitrogen | A21428 |
| Goat Anti-Mouse IgG (H+L), HRP | Jackson ImmunoResearch | 115-035-003 |
| Goat Anti-Rabbit IgG (H+L), HRP | Jackson ImmunoResearch | 111-035-003 |
| Mouse Anti-Goat IgG (H+L), HRP | Santa Cruz Biotechnology | sc-2354 |

(**M**): Monoclonal; (**P**): Polyclonal

### Supplementary Table S2. Primers and siRNAs used

| Primers used for mRNA gene expression | | |
|---|---|---|
| **Gene** | **Primer direction** | **Primer sequence** |
| AIRAP | Forward | 5′-TCATTTTCCATACGCTGCAC-3′ |
| | Reverse | 5′-CTGTGGTCCAAAGGGTGTCT-3′ |
| COX-2 | Forward | 5′-TTGCTGGCAGGGTTGCTGGTGGTA-3′ |
| | Reverse | 5′-CATCTGCCTGCTCTGGTCAATGGAA-3′ |
| HSP701A | Forward | 5′-CTACAAGGGGGAGACCAAGG-3′ |
| | Reverse | 5′-TTCACCAGCCTGTTGTCAAA-3′ |
| NKRF | Forward | 5′-CCAAACCTTCCAAAGGTCAA-3′ |
| | Reverse | 5′-CAGGGTTCCCACTGTCAAAA-3′ |
| L34 | Forward | 5′-GGCCCTGCTGACATGTTTCTT-3′ |
| | Reverse | 5′-GTCCCGAACCCCTGGTAATAGA-3′ |

| siRNAs sequences | |
|---|---|
| **siRNA** | **Target sequence** |
| **siHSF1$_1$** | 5′-TACCCAAGTACTTCAAGCACA-3′ |
| **siHSF1$_2$** | 5′-CAGTGACCACTTGGATGCTAT-3′ |

### Supplementary Figure Legends

**Figure S1. Induction of HSF1 phosphorylation by HCoV-229E at low multiplicity of infection.**

(**A**) Schematic representation of the experimental protocol. (**B**) Whole cell extracts from MRC-5 cells mock-infected (Mock) or infected with HCoV-229E (0.1 $TCID_{50}$/cell) were analyzed for pHSF1-Ser326, viral nucleocapsid (N) and α-tubulin protein levels at different times post infection (p.i.) by immunoblot.

**Figure S2. HCoV-induced expression of HSF1-target genes is dependent on the virus m.o.i. and persists at late times post infection.**

(A) Total mRNA was extracted from MRC-5 cells mock-infected or infected with HCoV-229E (0.1 or 1 $TCID_{50}$/cell) for 24h and analyzed for viral membrane (M-229E), HSP70 and AIRAP expression by qRT-PCR. (B) MRC-5 cells were mock-infected or infected with HCoV-229E (1 $TCID_{50}$/cell) and levels of M-229E, HSP70 and AIRAP mRNA were analyzed at different times p.i. by qRT-PCR. The fold increase was calculated by comparing the induction of M-229E, HSP70 and AIRAP in each sample to the mock-infected control (A) or to the relative control at 0h p.i. (B), which were arbitrarily set to 1. Error bars indicate means ± S.D.. * = $p < 0.05$; ANOVA test (A); Student's *t*-test (B).

**Figure S3. HCoV-OC43 and HCoV-NL63 infection induces the expression of HSF1-target genes in human cells.**

(**A, B**) Schematic representation of the experimental protocol (top panels). Immunoblot analysis of HSF1, HSPA6, AIRAP, viral N, α-tubulin and β-actin protein levels in MRC-5 (A) and Caco-2 hACE2 (B) cells mock-infected (-) or infected (+) with HCoV-OC43 (A) for 24h and HCoV-NL63 (B) for 72h at a m.o.i. of 0.1 $TCID_{50}$/cell (bottom panels).

**Figure S4. HSF1 activation is required for efficient replication of HCoV-OC43 coronavirus in different types of human cells.**

(**A**) Wild type (wt) or stably HSF1-silenced (HSF1i) HeLa cells were co-transfected with HCoV-OC43 genomic RNA (OC43 gRNA) and the pCMV-GFP vector for 4h. After 72h, levels of HSF1, viral spike (S) and nucleocapsid (N), GFP and GAPDH proteins were analyzed by IB (top panels). Relative amounts of S and N proteins were determined after normalizing to GAPDH (bottom panels). (**B**) Virus yield in the supernatant of cells treated as in A was determined by $TCID_{50}$ infectivity assay. Data, expressed as $TCID_{50}$/ml, represent the mean ± S.D. of duplicate samples. * = $p < 0.05$; Student's *t*-test. (**C**) MRC-5 cells mock-infected or infected with HCoV-OC43 (0.1 $TCID_{50}$/cell) were treated with different concentrations of the HSF1 inhibitor DTHIB immediately after the adsorption period. Virus yield was determined at 24h p.i. by infectivity assay. Data, expressed as $TCID_{50}$/ml, represent the mean ± S.D. of duplicate samples. * = $p < 0.05$.

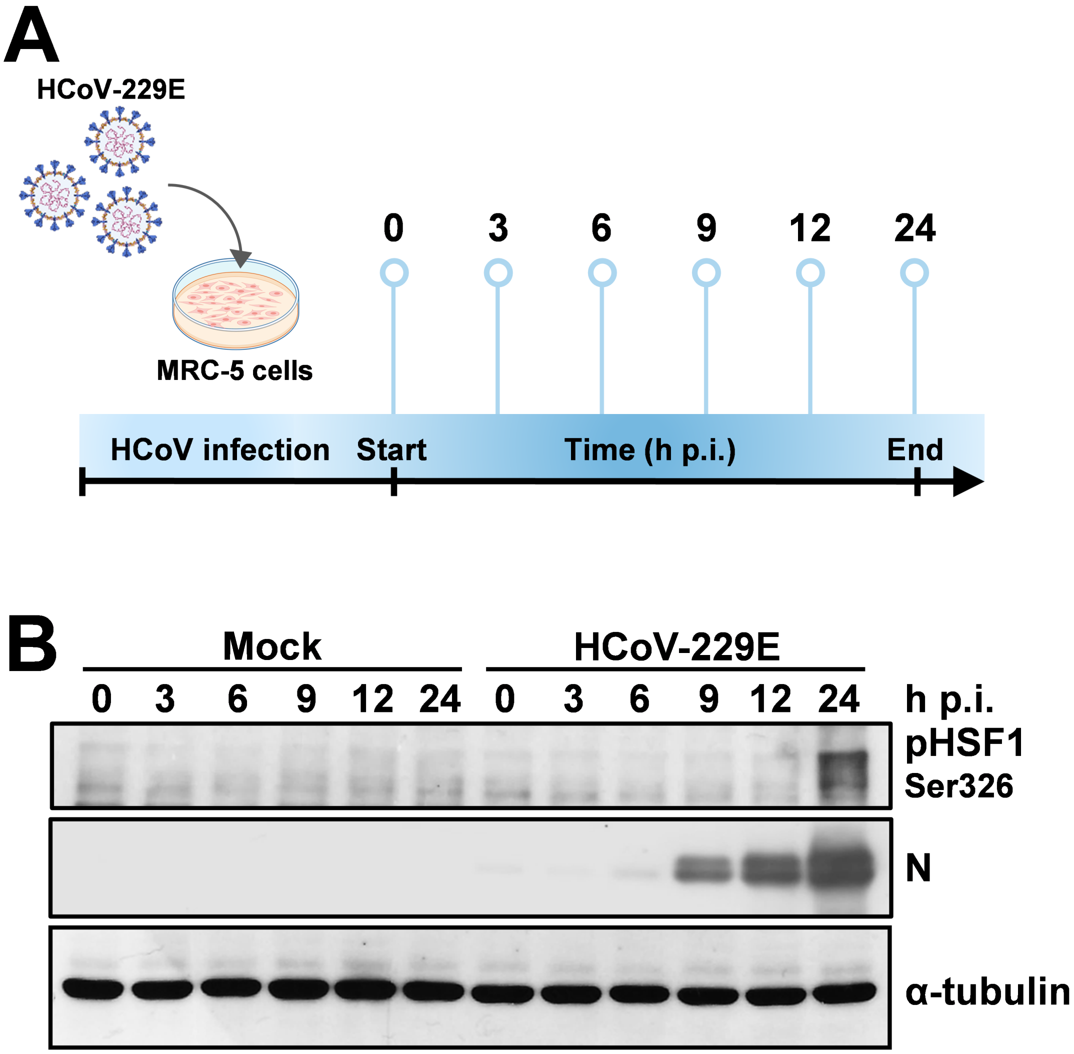
A
HCoV-229E
MRC-5 cells
0
3
6
9
12
24
HCoV infection
Start
Time (h p.i.)
End
B
Mock
HCoV-229E
0 3 6 9 12 24 0 3 6 9 12 24
h p.i.
pHSF1
Ser326
N
α-tubulin

Figure S1

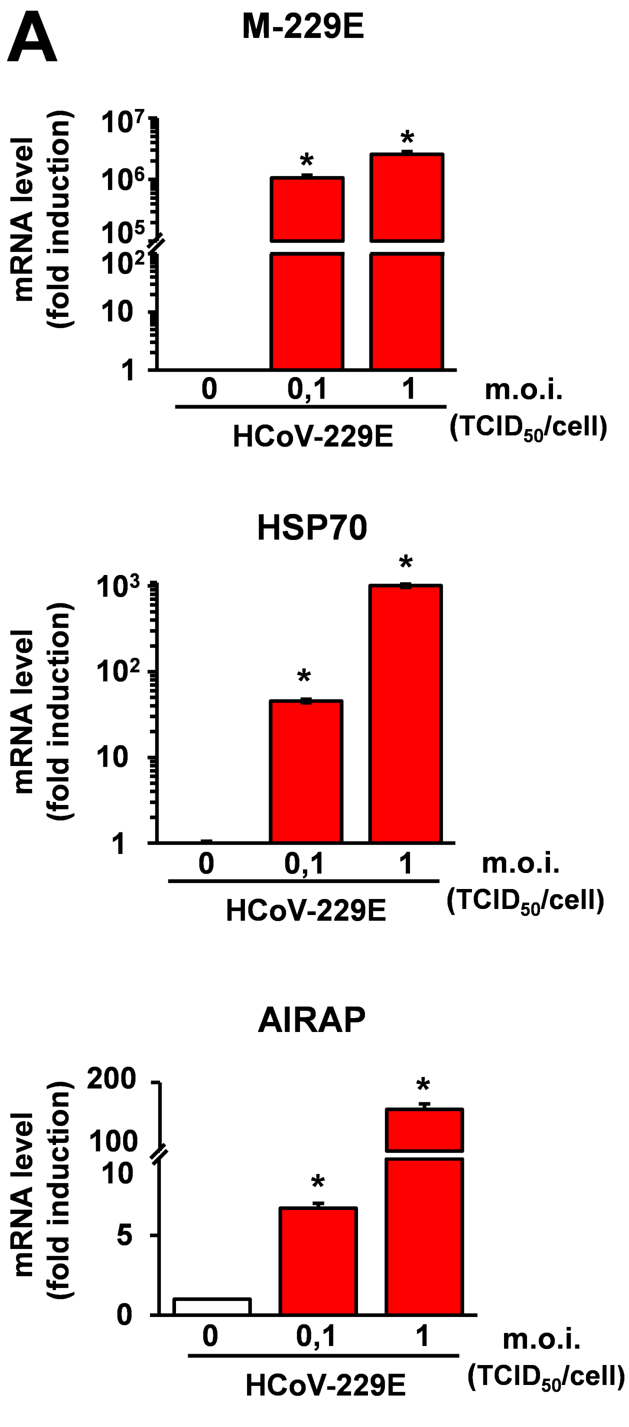
A
M-229E
mRNA level (fold induction)
0
0,1
1
m.o.i. (TCID50/cell)
HCoV-229E
HSP70
AIRAP
B
Mock
229E
Time (h p.i.)


Figure S2

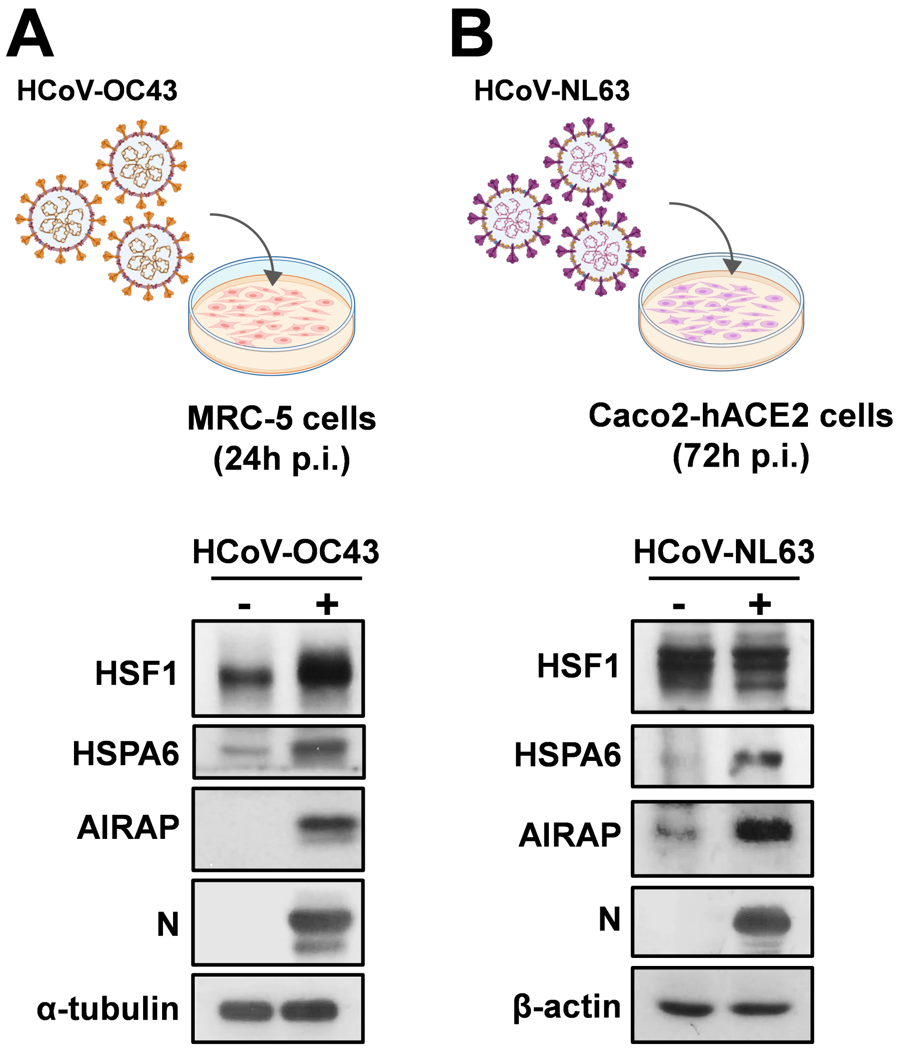
A
HCoV-OC43
MRC-5 cells
(24h p.i.)
HCoV-OC43
-
+
HSF1
HSPA6
AIRAP
N
α-tubulin
B
HCoV-NL63
Caco2-hACE2 cells
(72h p.i.)
HCoV-NL63
-
+
HSF1
HSPA6
AIRAP
N
β-actin

**Figure S3**

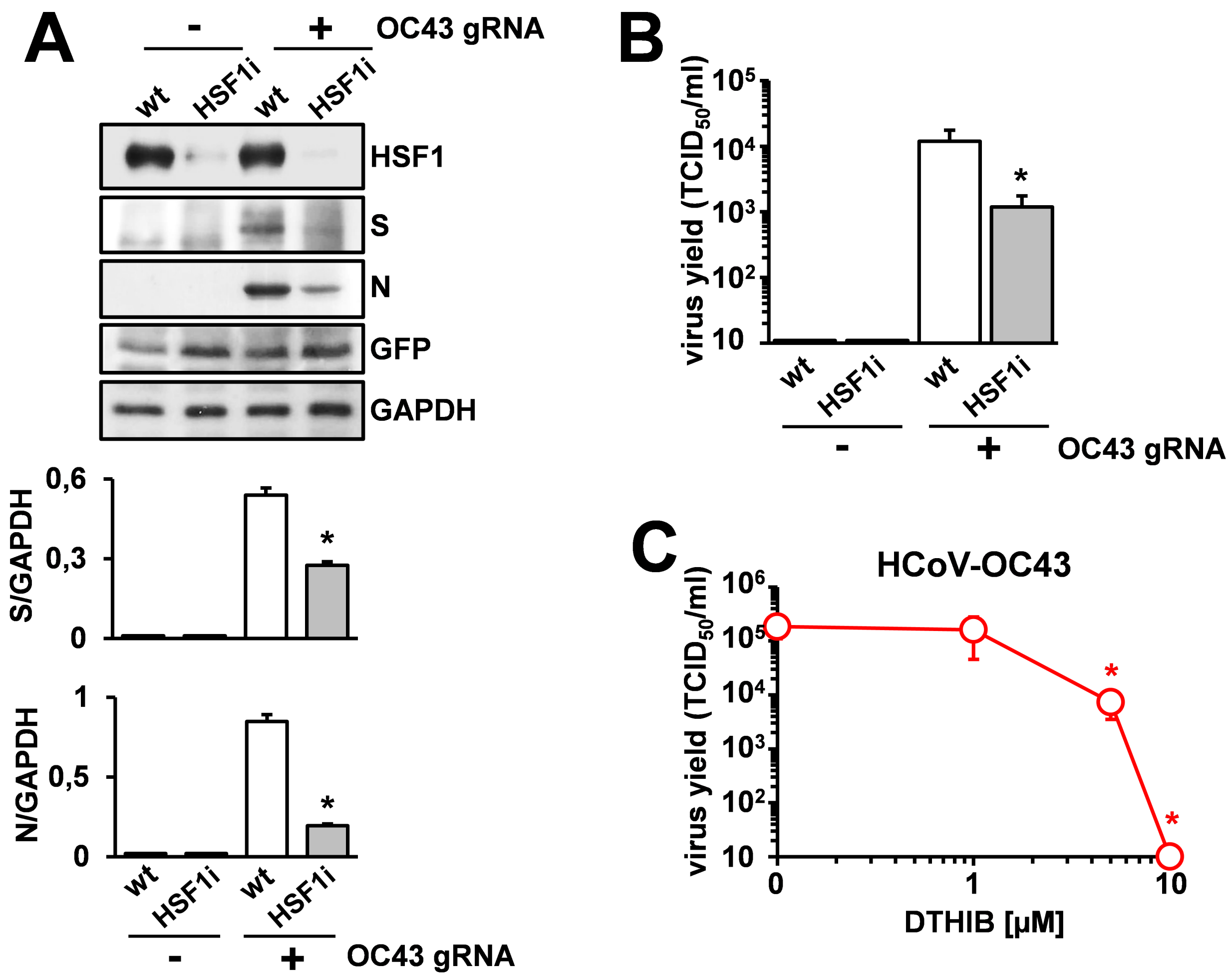
A
-
+
OC43 gRNA
wt
HSF1i
wt
HSF1i
HSF1
S
N
GFP
GAPDH
S/GAPDH
0,6
0,3
0
*
N/GAPDH
1
0,5
0
*
wt
HSF1i
wt
HSF1i
-
+
OC43 gRNA
B
virus yield (TCID50/ml)
10^5
10^4
10^3
10^2
10
*
wt
HSF1i
wt
HSF1i
-
+
OC43 gRNA
C
HCoV-OC43
virus yield (TCID50/ml)
10^6
10^5
10^4
10^3
10^2
10
*
*
0
1
10
DTHIB [µM]

Figure S4
